# Neural mechanisms of regulation of empathy and altruism by beliefs of others’ pain

**DOI:** 10.1101/2021.01.18.427136

**Authors:** Taoyu Wu, Shihui Han

**Author notes:** Address correspondence to: Shihui Han Ph.D. Prof. School of Psychological and Cognitive Sciences Peking University 52 Haidian Road, Beijing 100080, China.

## Abstract

Perceived cues signaling others’ pain induce empathy that in turn motivates altruistic behavior toward those who appear suffering. This perception-emotion-behavior reactivity is the core of human altruism but does not always occur in real life situations. Here, by integrating behavioral and multimodal neuroimaging measures, we investigate neural mechanisms underlying the functional role of beliefs of others’ pain in modulating empathy and altruism. We show evidence that decreasing (or enhancing) beliefs of others’ pain reduces (or increases) subjective estimation of others’ painful emotional states and monetary donations to those who show pain expressions. Moreover, decreasing beliefs of others’ pain attenuates neural responses to perceived cues signaling others’ pain within 200 ms after stimulus onset and modulate neural responses to others’ pain in the frontal cortices and temporoparietal junction. Our findings highlight beliefs of others’ pain as a fundamental cognitive basis of human empathy and altruism and unravel the intermediate neural architecture.

## Introduction

Aesop’s fable ‘The boy who cried wolf’ tells a story that villagers run or do not run to help a shepherd boy who cries wolf depending on whether they believe that the boy’s crying indicates his actual emotion and need. This story illustrates an important character of human altruistic behavior, that is, perceived cues signaling others’ suffering drives us to do them a favor only when we believe their suffering is true. Although this important character of human altruism was documented over 2000 years ago in Aesop’s fable and is widely observed in current human societies, its psychological and neural underpinnings have not been fully understood. The present study investigated how beliefs of others’ pain (BOP) modulate human altruistic behavior independently of perceived cues signaling others’ suffering and whether such modulation effects are mediated by changes in empathy for others’ pain and relevant brain underpinnings.

Empathy engages cognitive and affective processes that support understanding and sharing of other’s emotional states and provides a key motivation or a proximate mechanism for altruistic behavior in both humans and other animals (Batson et al., 2015; De Waal, 2008; Decety et al., 2016). Empathy can be induced by perceived cues signaling others’ pain that activate neural responses in brain regions underlying sensorimotor resonance (e.g., the sensorimotor cortex), affective sharing (e.g., the anterior cingulate cortex (ACC) and anterior insula (AI)), and mental state inference/perspective taking (e.g., the medial prefrontal cortex (mPFC) and temporoparietal junction (TPJ)) (Singer et al., 2004; Jackson et al., 2005; Avenanti et al., 2005; Fan and Han, 2008; Shamay-Tsoory et al., 2009; Han et al., 2009; Sheng and Han, 2012; Fan et al., 2011; Lamm et al., 2011; Zhou and Han, 2021). Neural responses to others’ pain in the empathy network and functional connectivity between its key hubs can predict motives of subsequent altruistic actions (Hein et al., 2010; 2016; Mathur et al., 2010; Luo et al., 2015). These findings revealed neural mechanisms underlying the perception(perceived pain)-emotion(empathy)-behavior(help) reactivity that occurs often in everyday lives (Eisenberg et al., 2010; Hoffman, 2008; Penner et al., 2005). However, empathic neural responses are influenced by multiple factors such as perceptual features depicting others’ pain (Gu and Han, 2007; Li and Han, 2019), observers’ perspectives and attention (Gu and Han, 2007; Li and Han, 2010; Jaunizux et al., 2019;), and perceived social relationships between observers and targets (Xu et al., 2009; Avenanti et al., 2010; Hein et al., 2010; Sheng and Han, 2012; Sheng et al., 2016; Han, 2018; Zhou and Han, 2021). What remains unclear is whether and how BOP modulates empathic brain activity by which to further influence altruistic behavior. To address these issues is crucial for understanding variations of empathy and altruism in social situations as that illustrated in the Aesop’s fable.

Beliefs refer to mental representations of something that is not immediately present to the scenes (Fuentes, 2019). As organism’s endorsement of a particular state of affairs as actual (McKay and Dennett, 2009), beliefs that best approximate reality enable the believers to act effectively and maximize their survival (Fodor, 1985; Millikan, 1995). Beliefs affect multiple mental processes such as visual awareness (Sterzer et al., 2008) and processing of emotions (Petrovic et al., 2005) including experiences of pain (Wager et al., 2004; Colloca and Benedetti, 2005). The function of beliefs is also manifested in increasing the efficiency of brain mechanisms involved in decision making and goal setting (Garces and Finkel 2019; Régner et al., 2019). Potential effects of beliefs on empathic neural responses were tested by presenting participants with photographs showing pain inflicted by needle injections into a hand that was believed to be or not to be anesthetized (Lamm et al., 2007). Functional magnetic resonance imaging (fMRI) of brain activity showed weak evidence for modulations of insular responses to perceived pain by beliefs of anesthetization. However, the results cannot be interpreted exclusively by BOP because the stimuli (i.e., needles) used to induce beliefs of numbed and non-numbed hands were different. An ideal paradigm for testing modulations of empathy by BOP independently of perceived cues signaling others’ pain should compare brain activities in response to identical stimuli under different beliefs and enable researchers to test how BOP influences altruistic behavior.

The current work tested the hypothesis that BOP provides a fundamental cognitive basis of empathy and altruistic behavior by modulating brain activity in response to others’ pain. Specifically, we tested predictions that weakening BOP inhibits altruistic behavior by decreasing empathy and its underlying brain activity whereas enhancing BOP may produce opposite effects on empathy and altruistic behavior. In Experiment 1, based on the common belief that patients show pain expressions to manifest their actual emotional state of pain whereas pain expressions performed by actors/actresses do not indicate their actual painful emotional states, we randomly assigned patient or actor/actress identities to faces with pain or neutral expressions to test how experimentally manipulated BOP changes associated with face identities cause changes in empathy (i.e., subjective evaluation of others’ pain) and altruistic behavior (i.e., monetary donations) in the directions as we predicted. In Experiment 2, based on the common belief that an effective medical treatment reduces a patient’s pain, we tested whether individuals’ intrinsic BOP predicts empathy and altruistic behavior across different faces with patient identities. In Experiments 3 and 4 we investigated whether BOP modulates empathic brain activity and showed electroencephalography (EEG) evidence that actor/actress compared to patient identities of faces decreased empathic neural responses to pain expressions of these faces within 200 ms post-stimulus. In Experiment 5 we further revealed behavioral and EEG evidence that neural responses to pain expressions of faces mediate BOP effects on empathy and monetary donations. Finally, in Experiment 6, we showed fMRI evidence that BOP modulated blood oxygen level dependent (BOLD) signals in the cognitive nodes of the empathic neural network in response to perceived painful stimulations. Together, our behavioral and neuroimaging findings indicate BOP as a fundamental cognitive basis for human empathy and altruism and uncover intermediate brain mechanisms by which BOP influences empathy and altruistic behavior.

## Results

### Experiment 1: Manipulated BOP changes empathy and altruistic behavior

In Experiment 1 we tested the predictions that weakening BOP decreases empathy and altruistic behavior whereas enhancing BOP produces opposite effects. We tested these predictions by experimentally manipulating individuals’ BOP. We presented participants (N = 60) with photos of faces of 16 models (half males) with pain expressions (see Methods for details). The participants were informed that these photos were taken from patients who suffered from a disease. In the 1^st^_round test the participants were shown with each photo and asked to report perceived pain intensity of each patient by rating on a Likert-type scale (0 = not painful at all; 10 = extremely painful) to evaluate their understanding of others’ pain — a key component of empathy (Jackson et al., 2005). Thereafter, the participants were invited to donate money to the patient in the photo by selecting an amount from an extra bonus payment for their participation (0 to 10 points, 1 point = ¥0.2) as a measure of altruistic behavior. The participants were informed that the amount of one of their decisions would be selected randomly and endowed to a charity organization to help those who suffered from the same disease. After the 1^st^_round test the participants were asked to perform a 5-minute calculation task to clean their memory of performances during the 1^st^_round test. The participants were then informed that this experiment actually tested their ability to recognize facial expressions and the photos were actually taken from 8 patients and 8 actors/actresses. We expected that these manipulations would influence BOP in opposite directions, that is, identity changes from patients to actors/actresses decreased BOP because patients’ pain expressions reflect their actual emotional states whereas pain expressions performed by actors/actresses do not indicate an actual painful state. By contrast, reconfirming patient identities enhanced the coupling between perceived pain expressions of faces and the painful emotional states of face owners and thus increased BOP. The participants were then asked to perform the 2^nd^_round test in which each photo was presented with patient or actor/actress identity indicated by a word below the photo. The participants had to rate perceived pain intensity of each photo and to report how much they would like to donate from the extra bonus payment to each model. The participants were told that an amount of money would be finally selected from their 2^nd^_round donation decisions and presented to a charity organization after the study.

The mean rating scores of pain intensity and amounts of monetary donations were subject to repeated-measures analyses of variance (ANOVAs) of Test Phase (1^st^_round vs. 2^nd^_round test) × Belief Change (Decreased BOP (patient to actor/actress) vs. Enhanced BOP (patient to patient)) as independent within-subjects variables. As expected, the results revealed that decreasing or enhancing BOP produced opposite effects on both perceived pain intensity and amounts of monetary donations, as indicated by significant Test Phase × Identity Change interactions (F(1,59) = 123.476 and 60.638, ps < 0.001, η_p_^2^ = 0.677 and 0.507, 90% CI = (0.555, 0.747) and (0.351, 0.611), Fig. 1a and 1b). Specifically, decreasing BOP by changing face identities from patients to actors/actresses significantly reduced perceived pain intensity and amounts of monetary donations in the 2^nd^_round (vs. 1^st^_round) test (F(1,59) = 82.664 and 34.542, ps < 0.001, η_p_^2^ = 0.584 and 0.369, 90% CI = (0.440, 0.673) and (0.207, 0.495)). By contrast, enhancing BOP by further confirming patient identities of faces significantly increased both perceived pain intensity and monetary donation in the 2^nd^_round (vs. 1^st^_round) test (F(1,59) = 36.060 and 27.457, ps < 0.001, η_p_ = 0.379 and 0.318, 90% CI = (0.216, 0.503) and (0.159, 0.449)). These results suggest that our manipulations of decreasing or enhancing BOP caused reliable changes in subjective evaluation of others’ pain and related monetary donations in opposite directions. Interestingly, to some degree rather than not at all, the participants reported pain and donated to faces with actor/actress identity in the 2^nd^_round test, suggesting that weakening BOP did not fully eliminate empathy and altruistic behavior toward those who showed pain expressions.

**Fig. 1.**
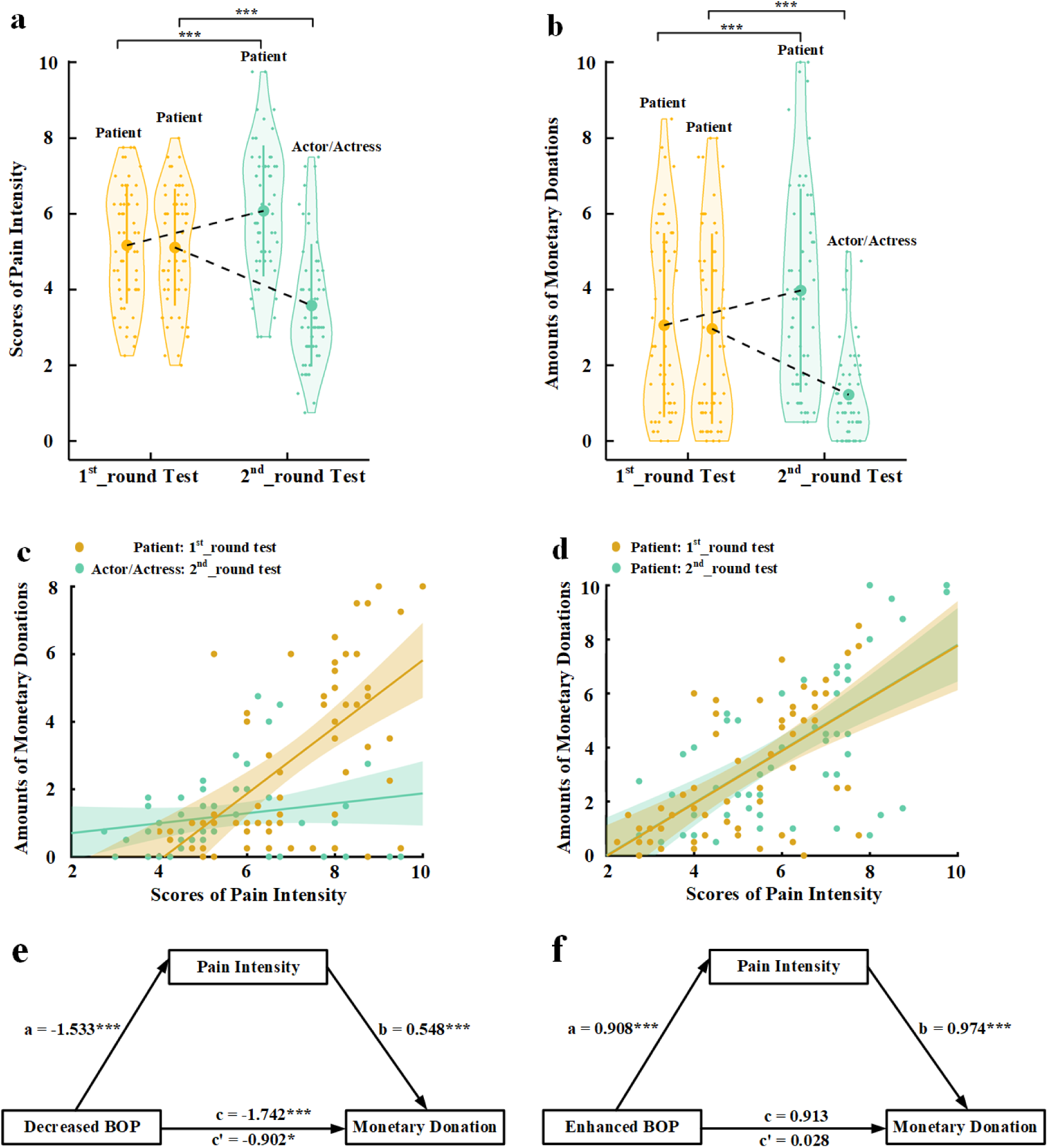
Behavioral results in Experiment 1. (a) Mean rating scores of pain intensity in the 1^st^_ and 2^nd^_round tests. (b) Mean amounts of monetary donations in the 1^st^_ and 2^nd^_round tests. Shown are group means (large dots), standard deviation (bars), measures of each individual participant (small dots), and distribution (violin shape) in (a) and (b). (c) The associations between rating scores of pain intensity and amounts of monetary donations for patients in the 1^st^_round test and for actors/actresses in the 2^nd^_round test. (d) The associations between rating scores of pain intensity and amounts of monetary donations for patients in both the 1^st^_ and 2^nd^_round tests. (e) Rating scores of pain intensity partially mediate the relationship between decreased BOP and reduced monetary donations. (f) Rating scores of pain intensity mediate the relationship between enhanced BOP and increased monetary donations.

To investigate whether perceived pain intensity mediated the relationships between experimentally manipulated BOP and monetary donation, we first conducted Pearson correlation analyses of the relationship between empathy and altruism. The results showed that the rating scores of pain intensity of faces whose identities changed from patient in the 1^st^_round test to actor/actress in the 2^nd^_round test significantly predicted the amount of monetary donations in the 1^st^_round but not in the 2^nd^_round test (r = 0.608 and 0.187, p < 0.001 and p = 0.152, 95% CI = (0.422, 0.776) and (−0.069, 0.435), all results were FDR-corrected, Fig. 1c). The rating scores of pain intensity also significantly predicted the amount of monetary donations for faces whose patient identities did not change in the 1^st^_round and 2^nd^_round tests (r = 0.619 and 0.628, ps < 0.001, 95% CI = (0.449, 0.776) and (0.417, 0.775), Fig. 1d). We conducted mediation analyses to further test an intermediate role of empathy between BOP and altruistic behavior (see Methods). The first mediation analysis showed that rating scores of pain intensity partially mediated the relationship between decreased BOP and reduced amount of monetary donations (direct effect: c’ = −0.902, t(118) = −2.468, p = 0.015, 95% CI = (−1.626, −0.178); indirect effect: a×b = −0.839, 95% CI = (−1.455, −0.374), Fig. 1e, see Supplementary Table 1_1 for statistical details). The second mediation analysis showed evidence that the rating scores of pain intensity also mediated the relationship between enhanced BOP and increased amount of monetary donations (direct effect: c’= 0.028, t(118) = 0.072, p = 0.943, 95% CI = (−0.727, 0.782), indirect effect: a×b = 0.885, 95% CI = (0.314, 1.563), Fig. 1f, see Supplementary Table 1_2 for statistical details). These results indicate a key functional role of BOP in altruistic behavior and suggest changes in subjective evaluation of others’ pain as an intermediate mechanism underlying the effect of BOP on monetary donations.

### Experiment 2: Intrinsic BOP predicts empathy and altruistic behavior

In Experiment 1 the experimenter manipulated BOP by assigning patient or actor/actress identities to faces with pain expressions and the results indicate that experimentally manipulated BOP changes caused variations of empathy and altruistic behavior. In Experiment 2 we further investigated whether an individual’s intrinsic BOP (i.e., various representations of actual emotional states of different faces with pain expressions) can predict his/her empathy and altruistic behavior across different faces. Moreover, as a replication, we tested whether changing the participants’ intrinsic BOP causes changes in empathy and altruistic behavior in a way similar to that observed in Experiment 1. In addition, we assessed whether changing intrinsic BOP modulated sharing of others’ pain — another key component of empathy (Jackson et al., 2005), and whether BOP induced emotional sharing mediates the relationship between BOP and altruistic behavior.

To address these issues, we tested an independent sample (N = 60) using the stimuli and procedure that were the same as those in Experiment 1 except the following. In the 1^st^_round test the participants were informed that they were to be shown with photos with pain expressions taken from patients who suffered from a disease and received a medical treatment. After the presentation of each photo the participants were asked to estimate, based on perceived pain expression of each face, how effective the medical treatment was for each patient by rating on a Likert-type scale (0 = no effect or 0% effective, 100 = fully effective or 100% effective). The rating scores were used to estimate the participants’ intrinsic BOP of each face with a higher rating score (indicating more effective treatment) corresponding to a weaker BOP as a more effective medical treatment reduces a patient’s pain to a greater degree. In addition to rating pain intensity of each face, the participants were asked to report how unpleasant they were feeling when viewing each photo by rating on a Likert-type scale (0 = not unpleasant at all, 10 = extremely unpleasant). The unpleasantness rating was performed to assess emotional sharing of others’ pain (Jackson et al., 2005). In the 2^nd^_round test the participants were told that the medical treatment was actually fully effective for half patients but had no effect for the others. Each photo was then presented again with information that the medical treatment applied to the patient was 100% effective (to decrease the participants’ beliefs of the patients’ painful states) or 0% effective (to enhance the participants’ beliefs of the patients’ painful states). Thereafter, the participants were asked to perform the rating tasks and to make monetary donation decisions, similar to those in the 1^st^_round test.

To assess whether individuals’ intrinsic BOP predicted their empathy and altruistic behavior across different target faces, we conducted Pearson correlation analyses of the relationships between intrinsic BOP as indexed by the rating score of treatment effectiveness and empathy rating scores/amounts of monetary donations across the sixteen models in the 1st_round test in each participant. The correlation coefficients were then transformed to Fisher’s z values that were further compared with zero. One-sample t-tests revealed that the z values were significantly smaller than zero (correlations between intrinsic BOP and pain intensity/unpleasantness/monetary donation: mean ± s.d. =-0.631 ± 0.531, −0.643 ± 0.524 and −0.469 ± 0.529; t(59) = −9.213, −9.501 and −6.875; ps < 0.001; Cohen’s d = 1.188, 1.227 and 0.887; 95% CI = (−0.768, −0.494), (−0.778, −0.507), and (−0.606, −0.333), Fig. 2a-c), suggesting that a larger score of treatment effectiveness (i.e., a weaker intrinsic BOP related to a face) predicted weaker empathy and less monetary donations relate to that face. These results provide evidence for associations between intrinsic BOP and empathy/altruism.

**Fig. 2.**
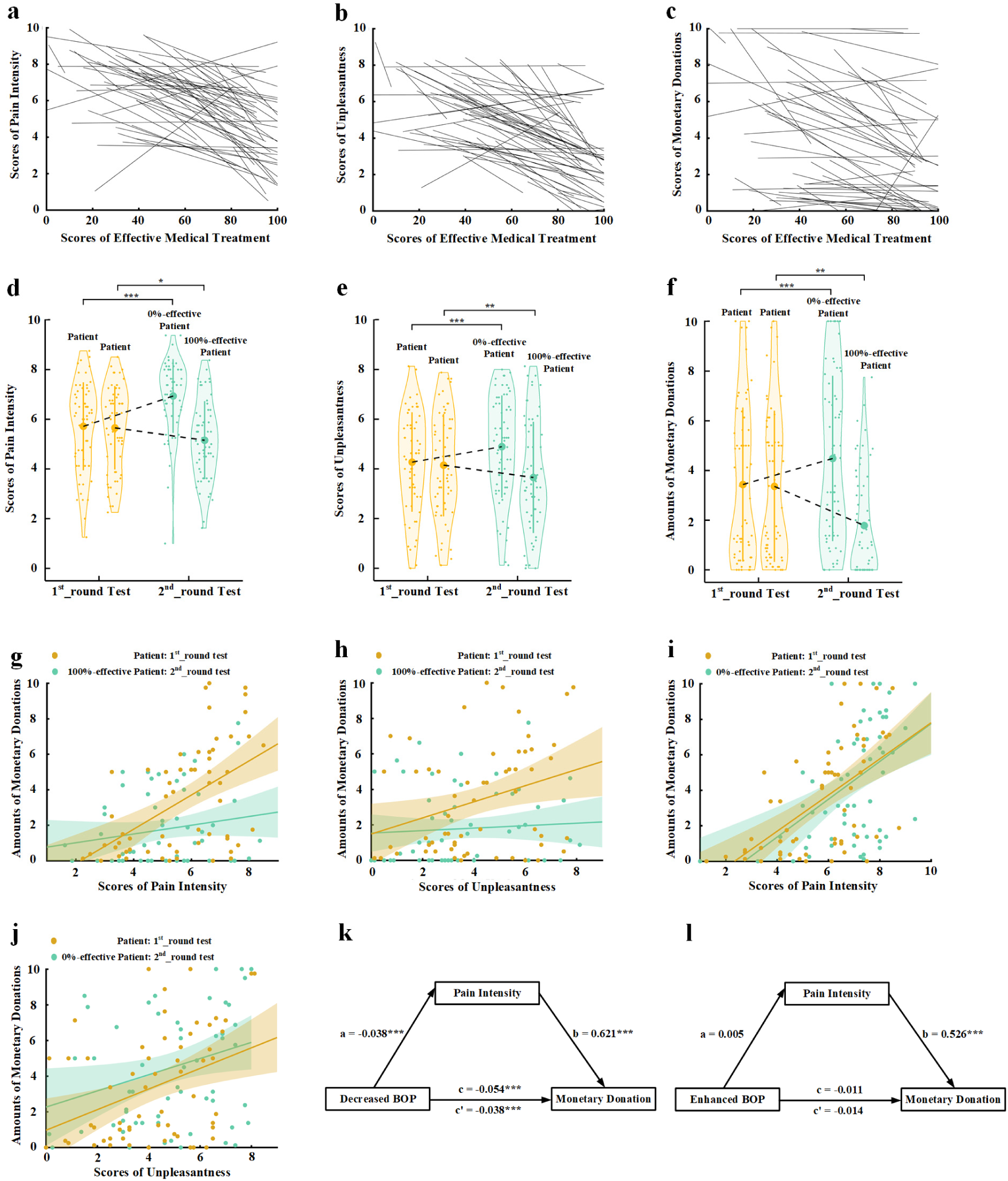
Behavioral results in Experiment 2. The relationships between intrinsic BOP (indexed by the rating score of effective medical treatments) and scores of pain intensity (a), own unpleasantness (b), and monetary donations (c), respectively, across the sixteen models in the 1st_round test in each participant. The regression line of each participant is plotted in (a), (b), and (c). (d-f) Mean rating scores of pain intensity, own unpleasantness, and monetary donations in the 1^st^_ and 2^nd^_round tests. (g) The associations between rating scores of pain intensity and amounts of monetary donations for patients in the 1^st^_round test and for 100%-effective patients in the 2^nd^_round tests across all the participants. (h) The associations between rating scores of own unpleasantness and amounts of monetary donations for patients in the 1^st^_round test and for-100% effective patients in the 2^nd^_round tests across all the participants. (i) The associations between rating scores of pain intensity and amounts of monetary donations for patients in the 1^st^_round test and for 0%-effective patients in the 2^nd^_round tests across all the participants. (j) The associations between rating scores of own unpleasantness and amounts of monetary donations for patients in the 1^st^_round test and for 0%-effective patients in the 2^nd^_round tests across all the participants. (k) Rating scores of pain intensity change partially mediate the relationship between decreased BOP and changes in monetary donations. (l) Rating scores of pain intensity change fail to mediate the relationship between enhanced BOP and changes in monetary donations. Shown are group means (large dots), standard deviation (bars), measures of each individual participant (small dots), and distribution (violin shape) in (d), (e), and (f).

Next, we tested whether decreased (or increased) BOP also predicts changes in empathy/altruistic behavior across different target faces for each participant. To do this, we calculated belief changes (decreased BOP: 100%-effective minus the participants’ initial estimation; enhanced BOP: the participants’ initial estimation minus 0%-effective), empathy changes (rating scores in the 2^nd^_round vs. 1^st^_round test), and changes in altruistic behavior (the amount of monetary donation in the 2^nd^_round vs. 1^st^_round test) related to each model in each participant. Similarly, we conducted Pearson correlation analyses to examine associations between changes in beliefs and empathy/donation for decreased-BOP patients and enhanced-BOP patients, respectively, in each participant. The correlation coefficients were then transformed to Fisher’s z values that were further compared with zero. One-sample t-tests showed that the z values were significantly smaller than zero for decreased-BOP patients (correlations between changes in belief and pain intensity/unpleasantness/monetary donation: mean ± s.d. = −0.304 ± 0.370, −0.277 ± 0.455 and −0.236 ± 0.410; t(59) = −6.352, −4.706 and −4.465; ps < 0.001; Cohen’s d = 0.822, 0.609 and 0.576; 95% CI = (−0.400, −0.208), (−0.394, −0.159), and (−0.342, −0.130)), as a greater decrease of BOP related to a face predicted greater reduced empathy and less monetary donations. By contrast, one-sample t-tests showed that the z values were significantly larger than zero for enhanced-BOP patients (correlations between changes in belief and pain intensity/unpleasantness/monetary donation: mean ± s.d. = 0.286 ± 0.488, 0.227 ± 0.470 and 0.162 ± 0.538; t(59) = 4.533, 3.735 and 2.332, ps < 0.001 and p = 0.023, Cohen’s d = 0.586, 0.483 and 0.301, 95% CI = (0.160, 0.412), (0.105, 0.348) and (0.023, 0.301)), as a greater increase of BOP related to a face predicted greater empathy and more monetary donations. These results provide evidence for associations between changes in BOP and empathy/altruism across different faces for each participant.

To test whether the results in Experiment 2 replicated those in Experiment 1, we conducted ANOVAs of the mean empathy scores and amounts of monetary donations with Test Phase (1^st^ vs. 2^nd^_round) and Belief Change (initial self-rated effectiveness to informed 0%-effectiveness vs. initial self-rated effectiveness to informed 100%-effectiveness) as independent within-subjects variables. The results showed that decreasing internal BOP (i.e., for 100% effective target faces) resulted in lower subjective evaluation of others’ pain and one’s own unpleasantness and less monetary donations in the 2^nd^_ vs. 1^st^_round tests, whereas enhancing BOP (i.e., for 0% effective target faces) produced opposite effects (Fig. 2d-f, see Supplementary Table 2_1 for statistical details). These results replicated those in Experiment 1 and provided further evidence that changing BOP resulted in variations of empathy and altruistic behavior.

Pearson correlations analyses of the mean rating scores in the 1^st^_round and 2^nd^_round tests across the participants showed that, for ‘100%-effective’ patients, the 1^st^_round but not the 2^nd^_round rating scores of empathy significantly predicted the amount of monetary donations (Pain intensity rating: r = 0.530 and 0.184, p < 0.001 and p = 0.159, 95% CI = (0.334, 0.698) and (−0.057, 0.425), Unpleasantness rating: r = 0.307 and 0.074, p = 0.017 and p = 0.576, 95% CI = (0.046, 0.541) and (−0.199, 0.358), Fig. 2g and 2h). For ‘0%-effective’ patients, however, both the 1^st^_round and 2^nd^_round rating scores of empathy significantly predicted the amount of monetary donations (Pain intensity rating: r = 0.582 and 0.476, ps < 0.001, 95% CI = (0.415, 0.725) and (0.287, 0.638); Unpleasantness rating: r = 0.373 and 0.280, p = 0.006 and 0.04, 95% CI = (0.096, 0.590) and (0.011, 0.511), Fig. 2i and 2j).

Furthermore, the results of mediation analyses showed that rating scores of pain intensity partially mediated the relationship between decreased BOP (i.e., for ‘100%-effective’ patients) and monetary donations ((direct effect: c’ = −0.038, t(58) = −3.657, p < 0.001, 95% CI = (−0.059, 0.017); indirect effect: a×b = −0.016, 95% CI = (−0.027, −0.005), Fig. 2k, see Supplementary Table 2_2 for statistical details). However, rating scores of unpleasantness did not mediate the relationship between decreased BOP and monetary donations (indirect effect: a×b = −0.002, 95% CI = (−0.009, 0.003)). Neither pain intensity nor unpleasantness ratings mediated the relationship between enhanced BOP (i.e., for ‘0%-effective’ patients) and monetary donations (indirect effect: a*b = 0.003 and −0.002, 95% CI = (−0.009, 0.013) and (−0.007, 0.004), Fig. 2l, see Supplementary Tables 2_3, 2-4, and 2-5 for statistical details). These results suggest that decreased BOP influences altruistic decisions possibly via modulations of the cognitive component of empathy (i.e., understanding others’ pain) rather than the affective component of empathy (i.e., sharing others’ pain).

### Experiment 3: Manipulated BOP changes empathic brain activity

Experiments 1 and 2 showed evidence that self-report measures of empathy for pain were affected by BOP. In Experiment 3 we further investigated whether and how changing BOP modulates brain activity in response to perceived cues signaling others’ pain as an objective estimation of empathy. If BOP provides a basis of empathy of others’ pain, decreasing BOP should reduce empathic neural responses to others’ pain. We tested this assumption by recording EEG to faces of 16 models from an independent sample (N = 30). The participants were first presented with these faces with neutral expressions and were informed that these photos were taken from 8 patients who suffered from a disease and from 8 actors/actresses. The participants were asked to remember patient or actor/actress identity of each neutral face and had to pass a memory test with a 100% recognition accuracy. Thereafter, the participants were informed that they would be presented with photos of these faces with either neutral or pain expressions. Photos of pain expressions were taken from the patients who were suffering from the disease whereas photos of pain expressions were taken from the actors/actresses who were performing pain expressions to imitate patients’ pain. The participants were asked to make judgments on identity of each face (i.e., patient vs. actor/actress) with a neutral or pain expression by pressing one of two buttons while EEG was recorded. After EEG recording, the participants were asked to rate pain intensity of each face with a pain or neutral expression on a Likert-type scale (0 = not painful at all; 7 = extremely painful) and to what degree they believed in the identity of each face with a pain expression on a 15-point Likert-type scale (−7 = extremely believed as an actor/actress, 0 = not sure, 7 = extremely believed as a patient). Because the same set of stimuli were perceived as patients or actors/actresses across the participants, modulations of brain activity in response to pain expressions only reflected the effects of BOP concomitant with the face identity (i.e., real pain for patients but fake pain for actors/actresses).

The participants reported a mean positive belief score corresponding to faces with a patient identity (2.496 ± 2.51) but a negative belief score corresponding to faces with an actors/actresses identity (−2.210 ± 3.25) (t(29) = 4.932, p < 0.001, Cohen’s d = 0.900, 95% CI = (2.755, 6.658)), suggesting successes of our manipulations of face identities. An ANOVA of the mean rating scores of pain intensity with Identity (patient vs. actor/actress) and Expression (pain vs. neutral) as within-subject variables revealed a significant Identity × Expression interaction (F(1,29) = 4.905, p = 0.035, η_p_^2^ = 0.145, 90% CI = (0.006, 0.330), Fig. 3a), suggesting greater subjective feelings of pain intensity for faces with patient compared to actor/actress identity. Moreover, a larger score of belief of patient identities significantly predicited greater subjective feeling of pain intensity related to patients’ pain (vs. neutral) expressions (r = 0.384, p = 0.036, 95% CI = (0.074, 0.627)), whereas there was no significant association between belief scores and subjective feeling of pain intensity related to actors/actresses’ pain (vs. neutral) expressions (r = 0.264, p = 0.159, 95% CI = (−0.162, 0.605)). These results provide further evidence for a link between BOP and empathy for patients’ pain.

**Fig. 3.**
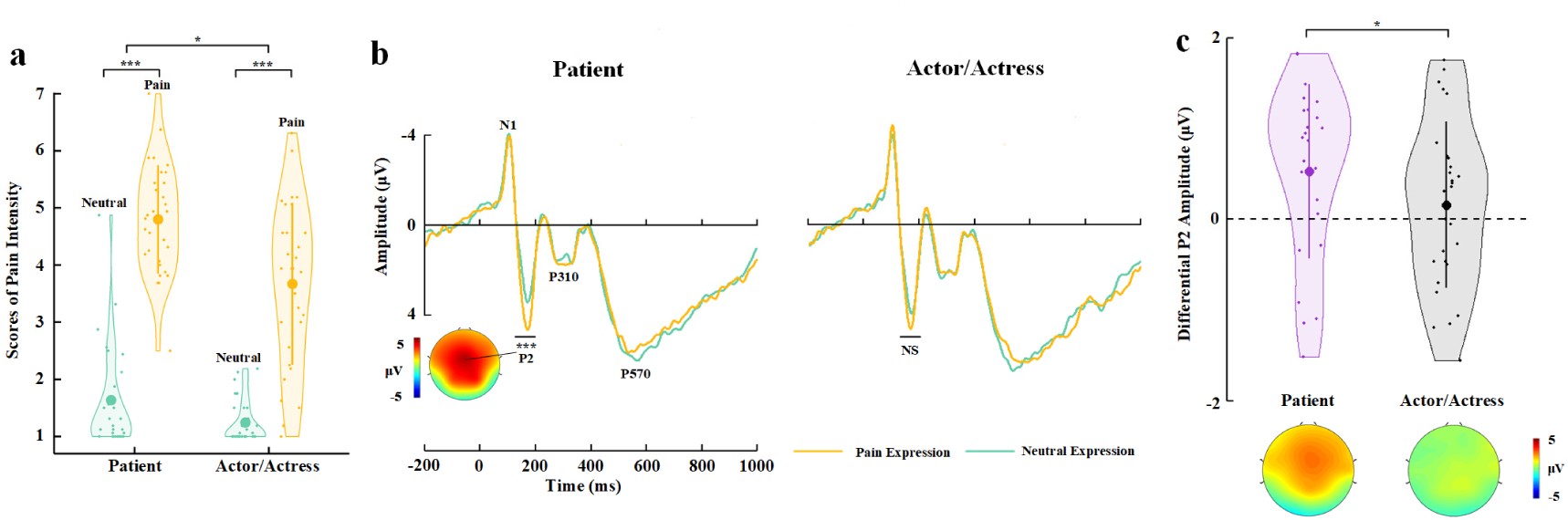
EEG results of Experiment 3. **(a)** Mean rating scores of pain intensity to pain vs. neutral expressions of faces with patient or actor/actress identities. (b) ERPs to faces with patient or actor/actress identities at frontal electrodes. The voltage topography shows the scalp distribution of the P2 amplitude with the maximum over the central/frontal region. (c) Mean differential P2 amplitudes to pain vs. neutral expressions of faces with patient or actor/actress identities. The voltage topographies illustrate the scalp distribution of the P2 difference waves to pain vs. neutral expressions of faces with patient or actor/actress identities, respectively. Shown are group means (large dots), standard deviation (bars), measures of each individual participant (small dots), and distribution (violin shape) in (a) and (c).

The participants responded to face identities with high accuracies during EEG recording (>81% across all conditions, see Supplementary Table 3_1 for details). ERPs to face stimuli in Experiment 3 were characterized by an early negative activity at 95-115 ms (N1) and a positive activity at 175-195 ms (P2) at the frontal/central regions, which were followed by two positive activities at 280-340 ms (P310) over the parietal region and 500–700 ms (P570) over the frontal area (Fig. 3b). Previous event-related potential (ERP) studies have shown that empathic neural responses to pain expressions are characterized by an increased P2 amplitude and the P2 amplitude to pain (vs. neutral) expressions predicts self-report of affective sharing (Sheng and Han, 2012; Sheng et al., 2016; Luo et al., 2018; Li and Han, 2019). Similarly, our ERP data analyses focused on whether BOP modulates the P2 amplitude to pain (vs. neutral) expressions given the previous ERP findings. ANOVAs of the P2 amplitudes with Identity (patient vs. actor/actress) and Expression (pain vs. neutral) as within-subject variables revealed a significant Identity × Expression interaction (F(1,29) = 7.490, p = 0.010, η_p_^2^ = 0.205, 90% CI = (0.029, 0.391), see Supplementary Table 3_2 for statistical details). Simple effect analyses verified significantly greater P2 amplitudes to pain vs. neutral expressions of patients’ faces (F(1,29) = 18.059, p < 0.001, η_p_ = 0.384, 90% CI = (0.150, 0.546)), whereas the P2 amplitude did not differ significantly between pain and neutral expressions of actors/actresses’ faces (F(1,29) = 0.334, p = 0.568, η_p_^2^ = 0.011, 90% CI = (0.000, 0.135), Fig. 3b and 3c). We further conducted Bayes factor analyses to examine the null effect of pain expressions on the P2 amplitudes to actors/actresses’ faces. The Bayes factor, which represents the ratio of the likelihood of the data fitting under the alternative hypothesis versus the likelihood of fitting under the null hypothesis, was 0.227, providing further evidence for the null hypothesis. The results indicate that, while the effect of pain (vs. neutral) expression on the P2 amplitudes to patients’ faces was similar to our previous findings that the P2 amplitudes increased to pain (vs. neutral) expressions of face without patient identities (Sheng and Han, 2012; Sheng et al., 2016), the P2 amplitude was less sensitive to pain vs. neutral expressions of faces with actor/actress identities. This finding provides evidence that decreasing BOP significantly weakens early empathic neural responses to others’ pain within 200 ms after stimulus onset.

### Experiment 4: BOP is necessary for modulations of empathic brain activity

The experimental manipulation in Experiment 3 consisted of multiple processes, including learning, memory, identity recognition, etc., to build different BOP related to different faces in the participants. If BOP rather than other processes was necessary for the modulation of empathic neural responses in Experiment 3, the same manipulation procedure to assign different face identities that do not change BOP should change the P2 amplitudes in response to pain expressions. We tested this prediction in an independent sample (N = 30) in Experiment 4. We employed the stimuli and procedure that were the same as those in Experiment 3 except that, during the learning phase, the participants were informed that the 16 models were from two baseball teams (half from a Tiger team and half from a Lion team) and they suffered from a disease. After the participants had remembered team identity of each neutral face in a procedure similar to that in Experiment 3, they performed identity (i.e., Tiger vs. Lion team) judgments on the faces with neutral or pain expressions during EEG recording. This manipulation built team identities should not influence self-report and EEG estimation of empathy because the Tiger/Lion team identities did not bring any difference in BOP between pain expressions of faces from the two teams.

The participants responded to face identities with high accuracies during EEG recording (> 79% across all conditions). Rating scores of pain intensity did not differ significantly between faces from the two teams (F(1,29) = 1.608, p = 0.215, η_p_^2^ = 0.053, 90% CI = (0, 0.216), Bayes factors = 0.261, Fig. 4a, see Supplementary Table 4_1 for details). ANOVAs of the mean P2 amplitudes over the frontal electrodes revealed a significant main effect of facial expression (F(1,29) = 12.182, P = 0.002, η_p_ = 0.296, 90% CI = (0.081, 0.473), Fig. 4b and 4c, see Supplementary Table 4_2 for details), as the P2 amplitude was enlarged by pain compared to neutral expressions. However, this effect did not differ significantly between faces from the two teams (F(1,29) = 0.040, P = 0.843, η_p_^2^ = 0.001, 90% CI = (0, 0.053), Bayes factors = 0.258). The null interaction effect on either self-report of empathy and the P2 amplitudes to pain (vs. neutral) expressions in Experiment 4 was not simply due to an underpowered sample size because the same sample size in Experiment 3 revealed reliable BOP effects on self-report and EEG (i.e., the P2 amplitude) estimation of empathy. Together, the results in Experiments 3 and 4 suggest a key role of BOP, but not other cognitive processes involved in the experimental manipulations, in modulations of neural responses to others’ pain.

**Fig. 4.**
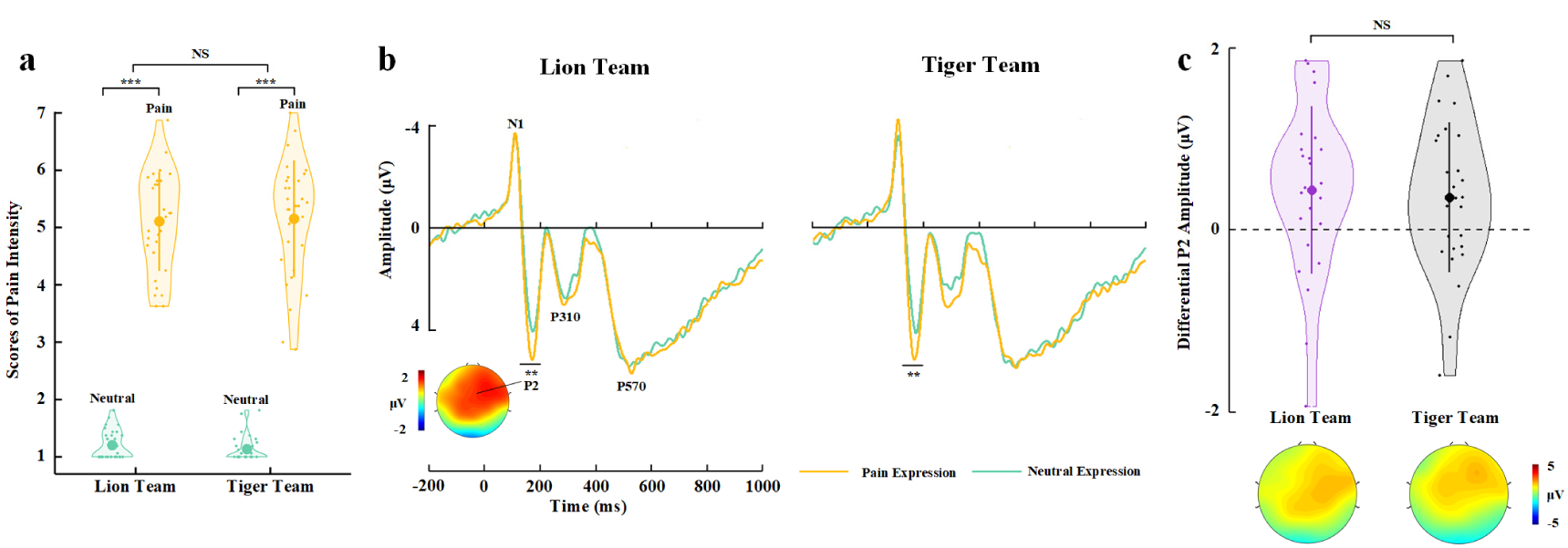
EEG results of Experiment 4. **(a)** Mean rating scores of pain intensity to pain vs. neutral expressions of faces with Lion Team or Tiger Team identities. (b) ERPs to faces with Lion/Tiger team identities at frontal electrodes. The voltage topography shows the scalp distribution of the P2 amplitude with the maximum over the central/frontal region. (c) Mean differential P2 amplitudes to pain vs. neutral expressions of faces with Lion/Tiger Team identities. The voltage topographies illustrate the scalp distribution of the P2 difference waves to pain vs. neutral expressions of faces with the Lion/Tiger Team identities, respectively. Shown are group means (large dots), standard deviation (bars), measures of each individual participant (small dots), and distribution (violin shape) in (a) and (c).

### Experiment 5: Empathic brain activity mediates relationships between BOP and empathy/altruistic behavior

Given that Experiments 1 to 4 showed consistent evidence for BOP effects on subjective feelings of others’ pain, altruistic behavior, and empathic neural responses, in Experiment 5, we further examined whether BOP-induced changes in empathic brain activity plays a mediator role in the pathway from belief changes to altered subjective feelings of others’ pain and altruistic decisions. To this end, we conducted two-session tests of an independent sample (N = 30). In the first session we employed the stimuli and procedure that were identical to those in Experiment 1 to assess BOP effects on empathy and altruistic behavior. In the second session we recorded EEG from the participants using the same stimuli and procedure as those in Experiment 3 to examine BOP effects on empathic neural responses. BOP-induced changes in empathic brain activity, rating scores of pain intensity, and amounts of monetary donations recorded in the two-session tests were then subject to mediation analyses.

To assure the participants’ beliefs about patient and actor/actress identities of perceived faces, after EEG recording, we asked the participants to complete an implicit association test (IAT) (Greenwald et al., 1998) that measured reaction times to faces with patient and actor/actress identities and words related to patients and actors/actresses (see Methods). The D score was then calculated based on response times (Greenwald et al., 2003) to assess implicit associations between patient and actor/actress faces and the relevant words. One-sample t-test revealed that the D score was significantly larger than zero (0.929 ± 0.418, t(29) = 12.178, p < 0.001, Cohen’s d = 2.223, 95% CI = (0.773, 1.085)), suggesting that patient faces were more strongly associated with patient relevant words whereas actor/actress faces were more strongly associated with actor/actress relevant words. The results indicate success of our belief manipulations through the two-session tests.

The behavioral results in the first-session test replicated the findings of Experiment 1. In particular, decreasing BOP (i.e., changing patient identity in the 1^st^_round test to actor/actress identity in the 2^nd^_round test) significantly reduced self-report of others’ pain and monetary donations (Test Phase × Identity Change interactions on rating scores of pain intensity and amounts of monetary donations: (F(1,29) = 59.654 and 129.696, ps < 0.001, η_p_^2^ = 0.673 and 0.817, 90% CI = (0.479, 0.764) and (0.694, 0.868); Effects of patient-to-actor/actress identity change on rating scores of pain intensity and amounts of monetary donations: F(1,29) = 58.196 and 180.022, ps < 0.001, η_p_^2^ = 0.667 and 0.861, 90% CI = (0.472, 0.760) and (0.765, 0.900), Fig. 5a and 5b). However, enhanced BOP (i.e., patient identity in the 1^st^_round test was further confirmed in the 2^nd^_round test) failed to significantly increase rating scores of pain intensity and amounts of monetary donations (F(1,29) = 0.016 and 0.209, p = 0.901 and 0.651, η_p_^2^ = 0.001 and 0.007, 90% CI = (0, 0.022) and (0, 0.119)), possibly due to ceiling effects of our measures in the participants (i.e., larger mean rating scores of pain intensity and mean amounts of monetary donations in the 1^st^_round test in Experiment 5 than in Experiment 1).

**Fig. 5.**
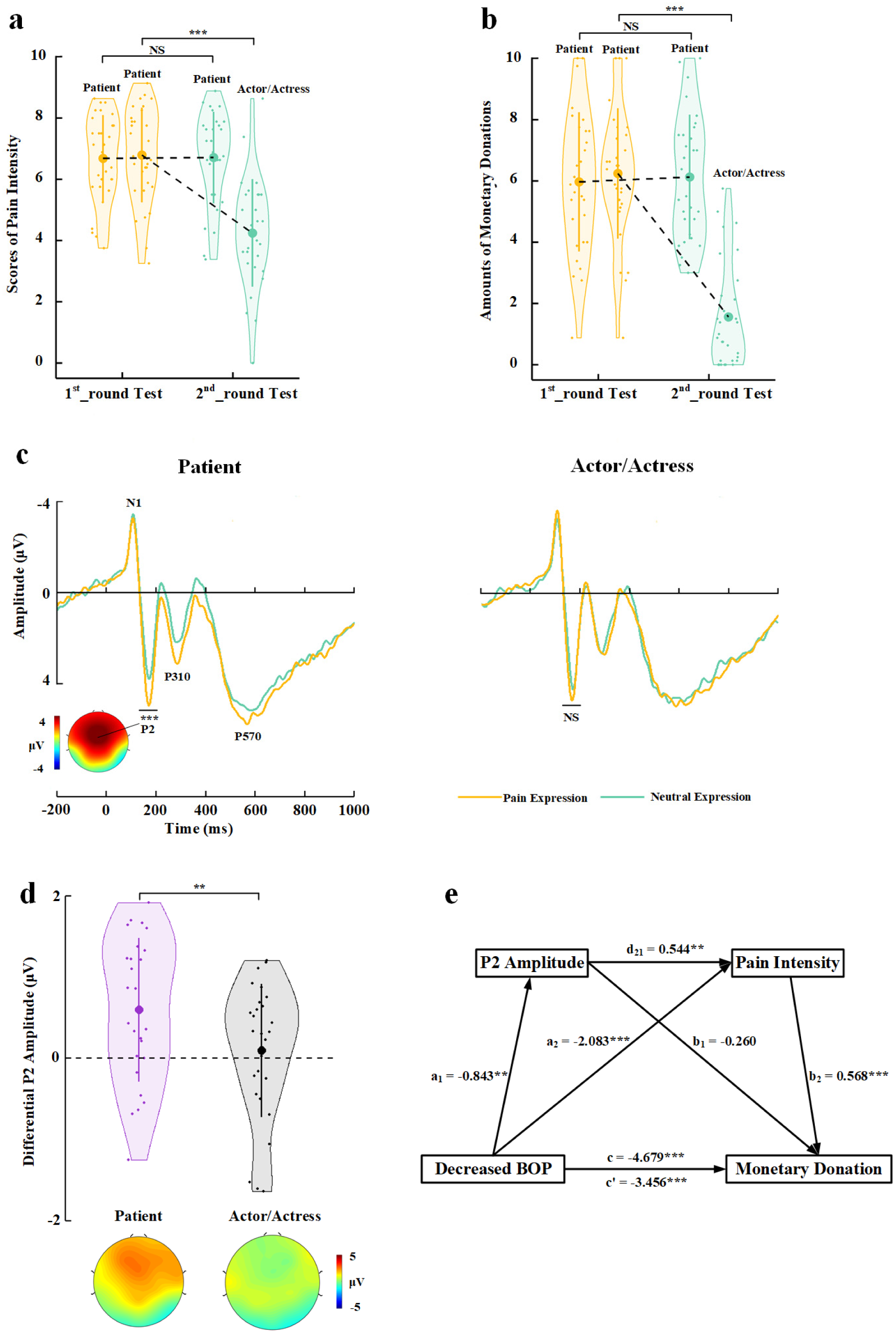
Behavioral and EEG results of Experiment 5. (a) Mean rating scores of pain intensity in the 1^st^_ and 2^nd^_round tests. (b) Mean amounts of monetary donations in the 1^st^_ and 2^nd^_round tests. (c) ERPs to faces with patient or actor/actress identities at frontal electrodes. The voltage topography shows the scalp distribution of the P2 amplitude with the maximum over the central/frontal region. (d) Mean differential P2 amplitudes to pain vs. neutral expressions of faces with patient or actor/actress identities. The voltage topographies illustrate the scalp distribution of the P2 difference waves to pain vs. neutral expressions of faces with patient or actor/actress identities, respectively. (e) Illustration of the serial mediation model of the relationship between decreased BOP and changes in monetary donations. Shown are group means (large dots), standard deviation (bars), measures of each individual participant (small dots), and distribution (violin shape) in (a), (b) and (d).

The participants responded to face identities with high accuracies during EEG recording (> 83% across all conditions). The EEG results replicated those in Experiment 3 by showing significantly deceased P2 amplitudes to pain (vs. neutral) expressions of actor/actress compared to patient faces (Identity × Expression interaction: F(1,29) = 9.494, p = 0.004, η_p_^2^ = 0.247, 90% CI = (0.050, 0.429), Fig. 5c and 5d, see Supplementary Table 5_1 for statistical details). Simple effect analyses verified significantly greater P2 amplitudes to pain vs. neutral expressions for patients’ faces (F(1,29) = 17.409, p < 0.001, η_p_^2^ = 0.375, 90% CI = (0.142, 0.539)) but not for faces of actors/actresses (F(1,29) = 0.270, p = 0.607, η_p_^2^ = 0.009, 90% CI = (0, 0.127), Bayes factor = 0.220). These behavioral and EEG results are consistent with those in Experiments 1 and 3 and provide repeated evidence for BOP effects on subjective feelings of others’ pain, altruistic behavior, and empathic brain activity in the same sample.

Next, we tested a serial mediation model of the relationship between decreased BOP (i.e., identity change from patient to actor/actress) and changes in monetary donations with two mediator variables including empathic neural responses (as indexed by the differential P2 amplitude to pain vs. neutral expressions) and changes in subjective feelings of others’ pain (as indexed by differential rating scores of pain intensity) (see Methods for details). This model includes three paths: (1) the indirect effect of BOP on monetary donation via the P2 amplitude (a_1_×b_1_ = 0.219, 95% CI = (−0.141, 0.745)); (2) indirect BOP effect on monetary donation via pain intensity (a_2_×b_2_ = −1.182, 95% CI = (−2.048, −0.510)); (3) indirect BOP effect on monetary donation via P2 amplitude × pain intensity (a_1_×d_21_×b_2_ = −0.261, 95% CI = (−0.584, −0.059), Fig. 5e, see Supplementary Table 5_2 for statistical details). The total indirect BOP effect on the monetary donation after controlling all indirect effect was c’ = −1.223, 95% CI = (−2.145, −0.400), which explained 26.14% variance of total effect of BOP on monetary donation. The effect sizes of the indirect path (2) and (3) were 25.26% and 5.58%, respectively, indicating that subjective feelings of others’ pain mediated the association between decreasing BOP and reduced monetary donation. Moreover, this mediator role was partially mediated by BOP induced variations of empathic brain activity in response to others’ pain expressions. Together, the results of these mediation analyses suggest a pathway from changes in BOP to varied empathic brain activity and changes in subjective report of empathy for other’s pain, which further accounted for BOP-induced changes in monetary donations.

### Experiment 6: Neural structures underlying BOP effects on empathy

While our EEG results revealed evidence for modulations of empathic neural responses by BOP, neural structures underlying the modulation effects remain unclear. In particular, it is unknown whether brain responses underlying cognitive and affective components of empathy are similarly sensitive to the influence of BOP. Therefore, in Experiment 6, we used fMRI to record BOLD signals from an independent sample (N = 31) to examine neural architectures in which empathic activities are modulated by BOP. Similarly, the participants were first shown with photos of neutral faces of 20 models and had to remember their patient (10 models) or actor/actress (10 models) identities. After the participants had performed 100% correct in a memory task to recognize the models’ identities, they were scanned using fMRI when viewing video clips of the models whose faces received painful (needle penetration) stimulation and showed pain expressions or received non-painful (cotton swab touch) stimulation and showed neutral expressions, similar to those used in the previous study (Han et al., 2009; Luo et al., 2014; Han et al., 2017). Before scanning the participants were informed that these video clips were recorded from the 10 patients who were receiving medical treatment and the 10 actors/actresses who practiced to imitate patients’ pain expressions. The participants responded to face identity (patient vs. actor/actress) of each model after viewing each video clip by pressing one of two buttons with high accuracies (> 80% across all conditions, see Supplementary Table 6_1 for details).

After fMRI scanning the participants were presented with each video clip again and had to rate the model’s pain intensity and their own unpleasantness. The participants were also asked to rate the degree to which they believed in the models’ patient or actor/actress identities in painful video clips on a 15-point Likert-type scale (−7 = extremely believed as an actor/actress, 0 = not sure, 7 = extremely believed as a patient) (see Method, Supplementary Table 6_1 for results). The mean rating scores confirmed significant differences in beliefs of patient and actors/actresses identities (2.776 ± 3.20 vs. −4.890 ± 1.44; t(30) = 10.526, p < 0.001, Cohen’s d = 1.890, 95% CI = (6.178, 9.153)), indicating success of our identity manipulations.

To localize empathic neural responses, we conducted whole-brain analyses of BOLD responses to perceived painful vs. non-painful stimuli applied to targets (collapsed faces with patient and actor/actress identities). This analysis revealed significant activations in the cognitive, affective, and sensorimotor nodes of the empathy network, including the bilateral insula/inferior frontal cortex (MNI peak coordinates x/y/z = −45/17/-5 and 45/26/-8), bilateral inferior and superior temporal gyri (−48/-70/-2 and 51/-58/-5), mPFC (3/56/25), left inferior parietal lobe (−63/-25/31), right superior parietal lobe (30/-58/55), and right post-central gyrus (58/-25/26, Fig. 6a; all activations were identified using a combined threshold of voxel level p < 0.001, uncorrected, and cluster level p < 0.05, FWE corrected). These brain activations are similar to those observed in previous research (Luo et al., 2014). To examine brain activity engaged in representing facial identities independent of perceived painful stimulation and pain expressions, we conducted a whole-brain analysis of the contrast of non-painful stimulations to patient vs. actor/actress. This analysis showed significant activations in the mPFC (−6/59/25) and bilateral TPJ (−54/-58/28 and 57/-67/31, Fig. 6b, all activations were identified using a combined threshold of voxel level p < 0.001, uncorrected, and cluster level p < 0.05, FWE corrected).

**Fig. 6.**
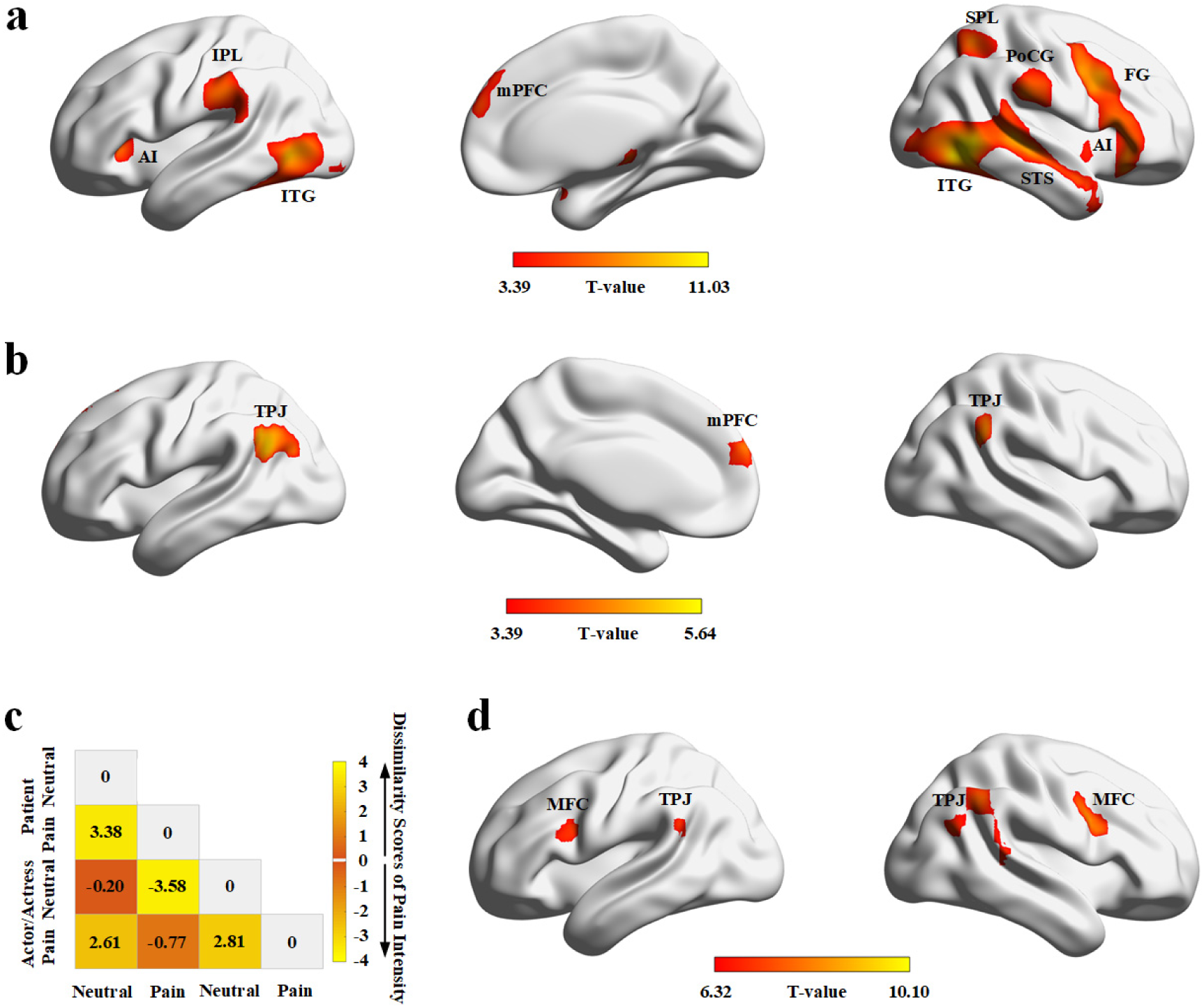
fMRI results of Experiment 6. (a) Brain activations in response to perceived painful (vs. non-painful) stimuli applied to targets (collapsed faces with patient and actor/actress identities). (b) Brain activations in response to non-painful stimuli to patients compared to actors/actresses. (c) Illustration of the behavioral dissimilarity matrix derived from the rating scores of pain intensity across all participants. Each cell in the dissimilarity matrix represents the mean difference in rating scores of pain intensity between each pair of conditions. (d) Brain activations that were correlated with the behavioral dissimilarity matrix revealed in the whole-brain searchlight RSA. AI = Anterior Insula; IPL = Inferior Parietal Lobe; ITG = Inferior Temporal Gyrus; mPFC = medial Prefrontal Cortex; SPL = Superior Parietal Lobe; PoCG = Post-Central Gyrus; FG = Frontal Gyrus; STS = Superior Temporal Sulcus; MFC = Middle Frontal Cortex; TPJ = Temporoparietal Junction.

Finally, we examined neural correlates of BOP effects on subjective feeling of others’ pain by conducting a representational similarity analysis (RSA) (Nili et al., 2014) of brain activity using a dissimilarity matrix (DM) constructed from scores of pain intensity in different conditions. To do this, we first conducted ANOVAs of the mean rating scores and found a significant Identity (patient vs. actor/actress) × Expression (pain vs. neutral) interaction on the rating scores of pain intensity (F(1,30) = 5.370, p = 0.027, η_p_^2^ = 0.152, 90% CI = (0.029, 0.391)) but not on the rating scores of unpleasantness (F(1,30) = 3.945, p = 0.056, η_p_^2^ = 0.116, 90% CI = (0, 0.296), see Supplementary Table 6_1 for statistical details). The results suggested a clear boundary between subjective feelings of pain intensity in different conditions. Next, we constructed a 4 × 4 DM for each participant with each cell in the DM representing the mean difference in rating scores of pain intensity between each pair of conditions, as illustrated in Fig. 6c. We then conducted a whole-brain searchlight RSA to identify brain regions in which the pairwise similarity of neural responses in the 4 conditions (2 Expressions × 2 Identities) corresponded to the behavioral DM in each participant (see Methods for details). The searchlight results were then subject to a second group-level analysis which revealed significant activations in the bilateral TPJ (−51/-46/26 and 54/-52/41) and bilateral middle frontal cortices (−33/2/26 and 36/-13/44, all activations were identified using RSA at a threshold of voxel-wise P (FWE) < 0.05, Fig. 6d). The results suggest that activities in the brain regions that have been shown to support self-other distinction/perspective taking (e.g., TPJ, Shamay-Tsoory et al., 2009; Lamm et al., 2019) and emotion regulation (e.g., lateral frontal cortices, Ochsner and Gross, 2005; Etkin et al., 2015) varied in correspondence to the distinct patterns of subjective feelings of pain intensity of faces with patient and actor/actress identities.

## Discussion

We conducted six experiments to investigate psychological and neural mechanisms underlying BOP impacts on empathy and altruistic behavior in humans. We manipulated individuals’ BOP by randomly assigning patient or actor/actress identities to faces and estimated individuals’ intrinsic BOP by asking the participants to estimate effectiveness of medical treatments related to different faces with pain expressions. We measured brain activity using EEG and fMRI to examine BOP effects on empathic neural responses with high temporal and spatial resolutions, respectively. Our behavioral and neuroimaging results revealed critical functional roles of BOP in modulations of the perception-emotion-behavior reactivity by showing how BOP predicted and affected empathy/empathic brain activity and monetary donations. Our findings provide evidence that BOP constitutes a fundamental cognitive basis for empathy and altruistic behavior in humans.

Experiments 1 and 2 showed behavioral evidence that manipulated changes in BOP caused subsequent variations of empathy and altruism along the directions as we expected. Specifically, decreasing BOP concomitant with changes in others’ identities (from patient to actor/actress) or in effective medical treatment (from suffering due to a disease to recovery due to medical treatment) significantly reduced self-report of both cognitive (perceived intensity of others’ pain) and affective (own unpleasantness induced by perceived pain in others) components of empathy. Decreasing BOP also inhibited following altruistic behaviors that were quantified by the amount of monetary donations to those who showed pain expressions. By contrast, enhancing BOP by reassuring patient identities or by noting the failure of medical treatment related to target faces magnified both cognitive and affective aspects of empathy and prompted more monetary donations to target faces. In consistent with the effects of manipulated BOP on empathy and altruism across the participants, the results of Experiment 2 showed that individuals’ intrinsic BOP related to each target face predicted their self-report of empathy and altruistic behavior across different target faces. Moreover, decreased (or increased) intrinsic BOP also predicted changes in empathy/altruistic behavior across different target faces. These converging behavioral findings across different participants and across different target faces provide evidence for causal relationships between BOP and empathy/altruism.

Our results also showed that self-report of other’s pain intensity and own unpleasantness positively predicted altruistic behavior. Previous research using questionnaire measures has shown that cognitive and affective components of trait empathy are positively correlated with the amount of money shared with others in economic games (Edele et al., 2013; Li et al., 2019). These findings together support the posit that empathy, as either an instant emotional response to others’ suffering or a sustained personality trait of understanding and sharing others’ pain, is one of the key motivating factors for altruistic behavior (Batson, 1987; Batson et al., 2015; Eisenberg et al., 2010; Hoffman, 2008; Penner et al., 2005). Our mediation analyses of the behavioral data in both Experiments 1 and 2 further revealed that the effects of decreased BOP on monetary donations were mediated by self-report of others’ pain intensity but not of own unpleasantness. These results imply that the cognitive component of empathy may function as an intermediate mechanism of the BOP impact on altruistic behavior.

Our neuroimaging experiments went beyond behavioral estimation of the relationships between BOP and empathy/altruism by uncovering neural mechanisms underlying BOP effects on empathy for others’ pain. Experiments 3 and 5 repeatedly showed that frontal neural responses to pain (vs. neutral) expressions within 200 ms after face onset (indexed by the P2 amplitude over the frontal/central electrodes) were significantly reduced to faces with actor/actress identities compared to those with patient identities. The results in Experiments 3 and 4 indicate that BOP concomitant with face identity (i.e., patients’ pain expressions manifest their actual painful emotional states whereas actors/actresses’ pain expressions do not) rather than face identity (e.g., Tiger or Lion team identities) alone resulted in modulations of the P2 amplitudes to pain expressions in the direction that we expected. Numerous EEG studies have shown that the frontal P2 component responds with enlarged amplitudes to various facial expressions such as fear, anger, happy (Williams et al. 2006; Luo et al. 2010; Calvo et al. 2013) and pain (Sheng and Han, 2012; Sheng et al., 2013; 2016) expressions compared to neutral faces, suggesting early affective processing by differentiating emotional and non-emotional expressions. ERPs to others’ pain within 200 ms post-stimulus occur regardless of task demands and are associated with spontaneous empathy for pain (Fan and Han, 2008). Our ERP results indicate that BOP provides a cognitive basis for early spontaneous neural responses to others’ pain expressions. Moreover, the results in Experiment 5 showed that the early spontaneous empathic neural responses in the P2 time window mediated BOP effect on self-report of perceived pain intensity in others, which further mediated the relationship between the P2 empathic responses and the amount of monetary donations. These results highlight both early spontaneous neural responses to others’ pain and subjective feelings of others’ pain as intermediate mechanisms by which BOP influences altruistic behavior.

To identify neural architectures underlying BOP effects on empathy, we recorded BOLD responses, using fMRI, to perceived painful and non-painful stimuli applied to individuals with patient or actor/actress identities in Experiment 6. We showed that, while the contrast of perceived painful (vs. non-painful) stimulations activated the sensory (i.e., post-central gyrus), affective (i.e., insula), and cognitive (mPFC) nodes of the empathy network (Singer et al., 2004; Jackson et al., 2005; Shamay-Tsoory et al., 2009; Han et al., 2009; Fan et al., 2011; Lamm et al., 2011; Zhou and Han, 2021; Luo et al., 2014), patient compared to actor/actress identities were associated with increased activity in the mPFC and bilateral TPJ. More importantly, the results of searchlight RSA revealed that variations of TPJ and lateral frontal activities in response to painful/non-painful stimuli corresponded with the patterns of self-report of empathy for patients and actors/actresses’ pain. The mPFC and TPJ are the key nodes of the social brain network which support inference of others’ mental states (Adolphs, 2003; Blakemore, 2008; Lieberman, 2010;). These brain regions also constitute the neural network underlying cognitive empathy — the processes of taking others’ perspectives, mentalizing, and self-other distinction (Shamay-Tsoory, 2011; Lamm et al., 2019). Although our fMRI results showed activations in the bilateral AI in response to others’ suffering, BOP seemed to mainly modulate the mPFC and TPJ activities. These fMRI results imply that BOP may influence altruistic behavior through its modulations of cognitive empathy. Consistent with the fMRI results, the results of mediation analyses in Experiments 1, 2, and 5 showed that BOP effects on altruistic behavior were mediated by cognitive (evaluation of others’ pain intensity) but not affective (sharing others’ emotional states) components of empathy. The behavioral and neuroimaging findings together suggest the cognitive component of empathy and the underlying neural circuits as intermediate mechanisms by which BOP influences altruistic behavior.

Our behavioral and neuroimaging findings have implications for how we understand the general functional role of beliefs in social cognition and interaction. Empathy is supposed to originate from an evolved adaptation to quickly and automatically respond to others’ emotional states during parental care that is necessary for offspring survival in humans and other species (De Waal, 2008; Decety, 2011). In most cases of interactions among family members (i.e., between parents and offspring or between siblings) perceived cues signaling pain in a person manifest his/her actual emotional states that urge help from other family members. Such life experiences may set up a default belief that perceived painful stimulation to others and their facial expressions reflect individuals’ actual emotional states. This default belief provides a fundamental cognitive basis of reflexive and automatic empathy and empathic brain activity that further generates autonomic and somatic responses, as suggested by the perception-action model of empathy (Preston and de Waal, 2002). Nevertheless, when social interactions expand beyond family members to non-kin members and even strangers, perceived pain expressions or painful stimuli applied to others may not always manifest others’ actual emotional states because perceived painful cues may be fake in some cases. BOP in such situations may function as cognitive gate-control to modulate neural responses to perceived pain in others. This is necessary for monitoring social interactions to determine whether to help or to coordinate with those who appear suffering. Our findings illustrate how the perception-emotion-behavior reactivity occurs under the cognitive constraint of BOP to keep empathy and altruistic decision/behavior for the right target who is really in need of help. In this sense, BOP also provides an important cognitive basis for survival and social adaption during complicated social interactions.

BOP effects on empathy and altruistic behavior can be understood in a framework of top-down modulations of empathic neural responses to others’ pain. Previous research has shown that, relative to the task of evaluating pain intensity of painful stimulation applied to others’ hands, counting the number of hands in stimulus displays (or paying less attention to painful cues) reduced neural responses in the ACC and AI (Gu and Han, 2007). Perceived information about social relationships between observers and those who appear suffering also modulates empathic neural responses to others’ pain such that, relative to viewing own-race or own-team individuals’ pain, viewing other-race or opponent-team individuals’ pain decreased empathic neural responses in the affective (e.g., ACC, AI), cognitive (e.g., mPFC, TPJ), and sensorimotor (e.g., motor cortex) nodes of the empathy network (Xu et al., 2009; Avenanti et al., 2010; Hein et al., 2010; Mathur et al., 2010; Sheng and Han, 2012; Sheng et al., 2014; 2016; Han, 2018; Zhou and Han, 2021). The perceived intergroup (racial) relationships between empathizers and the empathized also influenced altruistic behavior such as medical treatment (Drwecki et al., 2011). Although both BOP and perceived information about intergroup relationship influence altruistic behavior, the underlying neural mechanisms may be different since BOP effects on empathic neural responses seem to be limited to the cognitive empathic network whereas perceived intergroup relationships affect both the affective and cognitive components of the empathy network (Han, 2018).

It should be noted that our experimental manipulations changed the participants’ mind about the models’ identities (e.g., patient vs. actor/actress) rather than explicitly asking them to alter their BOP. BOP altered implicitly with target persons’ identities due to observers’ knowledge about individuals with different identities (e.g., painful stimuli applied to actors/actresses do not really hurt them and they show facial expressions to pretend a specific emotional state). Therefore, the BOP effects on empathy and altruistic behavior identified in our study might take place implicitly. This is different from the placebo effects on first-hand pain experiences that are produced by explicitly perceived verbal, conditioned, and observational cues that induce expectations of effective analgesic treatments (Meissner et al., 2011). Similar explicit manipulations of making individuals believe receiving oxytocin also promotes social trust and preference for close social distances (Yan et al., 2018). Moreover, the placebo treatment relative to a control condition significantly attenuated activations in the ACC, AI, and subcortical structures (e.g., the thalamus) in response to painful electric shocks but increased the prefrontal activity during anticipation of painful stimulations possibly to inhibit activity in pain processing regions (Wager et al., 2004; 2015). The brain regions in which empathic neural responses altered due to BOP (e.g., the mPFC and TPJ) as unraveled in the current study do not overlap with those in which activities are modulated by placebo analgesia (Atlas and Wager, 2014). These results suggest distinct neural underpinnings of BOP effects on empathic brain activity and placebo effects on brain responses to first-hand pain experiences.

Some limitations of the current work create future research opportunities. For example, a recent approach to hierarchical Bayesian models of cognition assumes that the brain represents information probabilistically and people represent a state or feature of the world not using a single computed value but a conditional probability density function (Knill and Pouget, 2004; Friston, 2005; Clark, 2013; Tappin and Gadsby, 2019). Our manipulations of BOP, however, had only two conditions (patient vs. actor/actress) and thus lack a model of effects of probability-based belief-updating on empathy and relevant altruistic behavior. Future research should examine how empathy and relevant altruistic behavior vary as a function of the degree of BOP. Other interesting research questions arising from our work include how the brain represents BOP. Different types of beliefs exist in human mind and may have distinct neural underpinnings (Harris et al., 2009; Seitz and Angel, 2020). To address neural representations of BOP will allow researchers to further explore and construct neural models of the interaction between beliefs and empathic brain activity in the key nodes of the empathy network. Another interesting issue related to our findings is individual differences in BOP and BOP effects on empathy and altruism. Since specific degrees of beliefs differ widely across individuals (Ais et al. 2016), it is crucial to examine what personality/psychopathic traits or biological factors make individuals hold strong or weak BOP and exhibit large or small BOP effects on empathy and altruistic behavior. It is also important to clarify what environmental factors modify individuals’ default BOP and consequently change their motivations to help those who appear suffering. To clarify these issues will advance our understanding of individual and contextual factors that shape the functional role of BOP in modulations of empathy and altruistic behavior. Finally, a general issue arising from the current work is whether beliefs affect the processing of other emotions such as fear, sad, and happy, and, if yes, whether there are common underlying psychological and neural mechanisms.

## Conclusion

Our behavioral and neuroimaging findings provide a new cognitive framework for understanding variations of human empathy and altruism. Our findings indicate that decreasing or increasing BOP produced opposite effects on empathy and altruistic behavior quantified by amounts of monetary donations. Changing BOP affected both subjective feelings of others’ emotional states and the underlying brain activity. BOP effects on altruistic behavior were mediated by two serial mediators, i.e., empathic neural responses and subjective feelings of others’ pain. Our behavioral and brain imaging findings together uncover BOP as a fundamental cognitive basis of the perception-emotion-behavior reactivity that underlies human altruism. The methods developed in our study open a new avenue for testing functional roles of beliefs as cognitive-gate control of other emotion processing and relevant social behavior.

## Methods

### Participants

Sixty Chinese students were recruited in Experiment 1 as paid volunteers (29 males, mean age ± s.d. = 21.15 ± 2.31 years). The sample size was estimated using G*Power (Faul et al., 2007) and a middle effect size of 0.25. To test the difference in pain intensity rating scores or monetary donations between the 1^st^_ and 2^nd^_round tests, we conducted ANOVA with Test Phase (1^st^ vs. 2^nd^_round) and Identity Change (patient to actor/actress vs. patient to patient) as independent within-subjects variables. To detect a significant Test x Identity interaction requires a sample size of 36 with an error probability of 0.05 and power of 0.95, given the correlation among repeated measures (0.5) and the nonsphericity correction (1). Sixty Chinese students were recruited in Experiment 2 as paid volunteers (30 males, 21.55 ± 2.45 years). Thirty Chinese students were recruited in Experiment 3 (all males, 22.23 ± 2.51 years) as paid volunteers. The sample size was determined based on our previous EEG research on empathy for pain using the same set of stimuli (Sheng and Han, 2012). We recruited only male participants to exclude potential effects of gender difference in empathic neural responses. Thirty-one Chinese students were recruited in Experiment 4 as paid volunteers. One participant was excluded from data analyses due to his lower response accuracy during EEG recording (< 50%). This left 30 participants (all males, 20.70 ± 1.97 years) for behavioral and EEG data analyses. Thirty Chinese students were recruited in Experiment 5 (all males, 20.60 ± 1.75 years). Thirty-two Chinese students were recruited in Experiment 6 as paid volunteers. One participant was excluded from data analyses due to excessive head movement during fMRI scanning. There were 31 participants left (all males, 22.23 ± 2.59 years) for behavioral and fMRI data analyses. The sample size in Experiment 6 was determined based on our previous fMRI research using similar stimuli (Luo et al., 2014). All participants had normal or corrected-to-normal vision and reported no history of neurological or psychiatric diagnoses. This study was approved by the local Research Ethics Committee of the School of Psychological and Cognitive Sciences, Peking University. All participants provided written informed consent after the experimental procedure had been fully explained. Participants were reminded of their right to withdraw at any time during the study.

### Experiment 1: Manipulated BOP changes empathy and altruistic behavior

#### Stimuli and procedure

The stimuli were adopted from our previous work (Sheng and Han, 2012), which consisted of photos of 16 Chinese models (half males) with each model contributing one photo with pain expression and one with neutral expression.

After reporting demographic information, the participants were informed that they would be paid with ¥10 as a basic payment for their participation. They would be able to obtain an extra bonus payment as much as ¥2 depending on their decisions in the following procedure. In the 1^st^_round test the participants were informed that they would be shown photos with pain expressions taken from patients who suffered from a serious disease. After the presentation of each photo the participants were asked to evaluate intensity of each patient’s pain based on his/her expression by rating on a Likert-type scale (0 = not painful at all; 10 = extremely painful). This rating task was adopted from previous research (Jackson et al., 2005) to assess the participants’ understanding of others’ pain feeling — a key component of empathy. Immediately after the pain intensity rating, the participants were asked to decide how much from the extra bonus payment they would like to donate to the patient (0 to 10 points, 1 point = ¥0.2). The participants were informed that the amount of one of their decisions would be selected randomly and donated to one of the patients.

After the 1^st^_round test the participants were asked to perform a short (5 mins) calculation task (10 arithmetic calculations, e.g. 25-3×7=?) to clean their memory of the 1^st^_round ratings. Thereafter, the participants were told that the photos were actually taken from 8 patients and 8 actors/actresses and this experiment actually tested their ability of recognizing social identities by examination of facial expressions. Faces assigned with patient or actor/actress identities were counterbalanced across the participants. The participants were then asked to conduct the 2^nd^_round test in which each photo was presented again with a word below to indicate patient or actor/actress identity of the face in the photo. The participants had to report again perceived pain intensity of each face and how much they would like to donate to the person shown in the photo. The participants were informed that an amount of money would be finally selected randomly from their 2^nd^_round decisions and donated to one of the patients. The total amount of the participants’ donations were subject to a charity organization to help patients who suffer from the same disease after the study.

We conducted ANOVAs of rating scores of pain intensity and amounts of monetary donations with Test Phase (1^st^ vs. 2^nd^_round) × Identity Change (patient to actor/actress vs. patient to patient) as independent within-subjects variables to assess whether and how beliefs of others’ pain (BOP) influenced empathy and altruistic behavior towards those who suffered. Finally, the participants completed two questionnaires to estimate individual differences in trait empathy (Davis, 1983) and interpersonal trust (Wright and Tedeschi, 1975). We analyzed the relationship between our empathy/altruistic measures and individuals’ trait empathy/interpersonal trust but failed to find significant results and thus were not reported in the main text.

#### Mediation analysis

We performed mediation analyses to examine whether pain intensity mediates the pathway from BOP to monetary donation. To do this, we first dummy coded decreased BOP as 0 (patient identity in the 1^st^_round test) and 1 (actor/actress in the 2^nd^_round test) or enhanced BOP as 0 (patient identity in the 1^st^_round test) and 1 (patient identity was confirmed in the 2^nd^_round test). Then, we estimated four regression models: 1) whether the independent variable (BOP) significantly accounts for the dependent variable (monetary donation) when not considering the mediator (e.g., Path c); 2) whether the independent variable (BOP) significantly accounts for the variance of the presumed mediator (pain intensity) (e.g., Path a); 3) whether the presumed mediator (pain intensity) significantly accounts for the variance of the dependent variable (monetary donation) when controlling the independent variable (BOP) (e.g., Path b); 4) whether the independent variable (BOP) significantly accounts for the variance of the dependent variable (monetary donation) when controlling the presumed mediator (pain intensity) (e.g., Path c’). To establish the mediation, the path c is not required to be significant. The only requirement is that the indirect effect a×b is significant. Given a significant indirect effect, if Path c is not significant, the mediation is classified as indirect-only mediation which is the strongest full mediation (Kenny et al., 1998; Zhao et al.., 2019). A bootstrapping method was used to estimate the mediation effect. Bootstrapping is a nonparametric approach to estimate effect-sizes and hypotheses of various analyses, including mediation (Shrout and Bolger,2002; Mackinnon et al., 2004). Rather than imposing questionable distributional assumptions, a bootstrapping analysis generates an empirical approximation of the sampling distribution of a statistic by repeated random resampling from the available data, which is then used to calculate p-values and construct confidence intervals. 5,000 resamples were taken for our analyses. Moreover, this procedure supplies superior confidence intervals (CIs) that are bias-corrected and accelerated (Preacher et al., 2007; Preacher and Hayes, 2008a, 2008b). The analyses were performed using Hayes’s PROCESS macro (Model 4, Hayes, 2017).

#### Statistical comparison

Behavioral data were assumed to have a normal distribution but this was not formally tested. 95% Confidence intervals (95% CIs) were reported for t-tests of the mean difference between two conditions and for correlation analyses of correlation coefficients. 90% CIs were reported for effect sizes (η_p_^2^) of ANOVA analyses. According to Steiger (2004), the general rule of thumb to use CIs to test a statistical hypothesis (H0) is to use a 100×(1-α)% / 100×(1-2α)% CI when testing a two-sided / one-sided hypothesis at alpha level. We thus reported 90% CIs of η^2^ in ANOVAs because η^2^ is always positive.

### Experiment 2: Intrinsic BOP predicts empathy and altruistic behavior

The face stimuli and the procedure were the same as those in Experiment 1 except the following. The participants were informed that they were to be shown photos with pain expressions taken from patients who had suffered from a serious disease and received medical treatment. After the presentation of each photo the participants were asked to estimate how effective the medical treatment was for each patient by rating on a Likert-type scale (0 = no effective or 0%, 100 = fully effective or 100% effective). Besides rating pain intensity of each face in the 1^st^_round test, the participants were asked to report how unpleasant they were feeling when they viewed the photo (i.e., own unpleasantness) by rating on a Likert-type scale (0 = not unpleasant at all, 10 = extremely unpleasant). The unpleasantness rating was performed to evaluate emotional sharing of others’ pain — another key component of empathy (Jackson et al., 2005). The order of the two empathy rating tasks was counterbalanced across the participants. Immediately after the empathy rating tasks, the participants were asked to decide how much from the extra bonus payment they would like to donate to the patient (0 to 10 points, 1 point = ¥0.2).

In the 2^nd^_round test the participants were told that the medical treatment was actually effective for only half of the patients. Each photo was then presented again with information that the medical treatment applied to the patient was 100% effective or 0% effective. Thereafter, the participants were asked to perform the rating tasks and monetary donation as those in the 1^st^_round test. The participants were told that an amount of money would be finally selected from their 2^nd^_round decisions and donated to one of the patients.

#### Mediation analysis

This was the same as that in Experiment 1 except that we tested whether changes of pain intensity mediate the pathway from decreased BOP or enhanced BOP to changes of monetary donation. To do this, we first calculated belief update (decreased BOP: 100%-effect minus the participants’ initial estimation; enhanced BOP: the participants’ initial estimation minus 0%(no)-effect). Then, we estimated four regression models: 1) whether the independent variable (BOP) significantly accounts for the dependent variable (changes of monetary donation) when not considering the mediator (e.g., Path c); 2) whether the independent variable (BOP) significantly accounts for the variance of the presumed mediator (changes of pain intensity) (e.g., Path a); 3) whether the presumed mediator (changes of pain intensity) significantly accounts for the variance of the dependent variable (changes of monetary donation) when controlling the independent variable (BOP) (e.g., Path b); 4) whether the independent variable (BOP) significantly accounts for the variance of the dependent variable (changes of monetary donation) when controlling the presumed mediator (changes of pain intensity) (e.g., Path c’).

### Experiment 3: Manipulated BOP changes empathic brain activity

#### Stimuli and procedure

Face stimuli were adopted from our previous work (Sheng and Han, 2012) and used in Experiments 3, 4 and 6 in this study. The stimuli consisted of 32 faces of 16 Chinese models (half males) with each model contributed one photo with pain expression and one with neutral expression. During behavioral tests or EEG recording, each photo was presented in the center of a gray background on a 21-inch color monitor, subtending a visual angle of 3.8° × 4.7° (width × height: 7.94 × 9.92 cm) at a viewing distance of 60 cm.

Before EEG recording the participants were asked to perform an identity memory task in which faces with neutral expressions were presented. Eight faces were marked as patients and 8 faces as actors/actresses. After viewing photos with marked identity for 15 minutes, the participants performed a discrimination task in which each neutral face was displayed for 200 ms and the participants had to press the left or right button using the left or right index finger to indicate identity of each face (i.e., patient or actor/actress) within two seconds. After their response accuracies reached 100%, the participants were moved into an acoustically- and electrically-shielded booth for EEG recording.

During EEG recording each trial consisted of a painful or neutral face with a duration of 200 ms, which was followed by a fixation cross with a duration varying randomly between 800 and 1400 ms. There were 8 blocks of 64 trials (each of the 32 photographs was presented twice in a random order in each block). The participants were asked to press the left or right button using the left or right index finger to indicate the identity of the face (i.e., patient or actor/actress) as fast and accurately as possible.

The relation between responding hand and face identity was counterbalanced across different blocks of trials.

After EEG recording, the participants were presented with each face again with a pain expression and asked to rate how painful the person is feeling (i.e., pain intensity) by rating on a Likert-type scale (1 = not painful at all; 7 = extremely painful). To estimate the participants’ BOP, they were also asked to answer the question of “To what extent do you believe the identity of this model (either patient or actor/actress)?” on a 15-point Likert-type scale (−7 = extremely believed as an actor/actress, 0 = not sure, 7 = extremely believed as a patient).

#### EEG data acquisition and analysis

A NeuroScan system (CURRY 7, Compumedics Neuroscan) was used for EEG recording and analysis. EEG was continuously recorded from 32 scalp electrodes and was re-referenced to the average of the left and right mastoid electrodes offline. Impedances of individual electrodes were kept below 5 kΩ. Eye blinks and vertical eye movements were monitored using electrodes located above and below the left eye. The horizontal electro-oculogram was recorded from electrodes placed 1.5-cm lateral to the left and right external canthi. The EEG trace was digitized at a sampling rate of 1,000 Hz and subjected to an online band-pass filter of 0.01–400 Hz. EEG data were filtered with a low-pass filter at 30 Hz offline. Artefacts related to eye movement or eye blinks were removed using the covariance analysis tool implemented in CURRY 7 (Semlitsch et al., 1986). Only trials with correct responses to face identity were included for data analyses (see Supplementary Table 7 for the numbers of trails included for data analyses in Experiments 3-5). The baseline for all ERP measurements was the mean voltage of a 200-ms prestimulus interval and the latency was measured relative to the stimulus onset.

Face stimuli in the identity judgment task elicited an early negative activity at 95-115 ms (N1) and a positive activity at 175-195 ms (P2), followed by a positive activity at 280-340 ms (P310) and a long-latency positivity at 500–700 ms (P570) over the frontal area. The mean ERP amplitudes were subject to ANOVAs with Identity (patient vs. actor/actress) and Expression (Pain vs. Neutral) as within-subject variables. To avoid potential significant but bogus effects on ERP amplitudes due to multiple comparisons (Luck and Gaspelin, 2017), the mean values of the amplitudes of the N1, P2, P310, and P570 components were calculated at frontocentral electrodes (i.e., F3, Fz, F4, FC3, FCZ and FC4).

To further assess the null hypothesis regarding the difference in the P2 amplitude in response to pain and neutral expressions of actors/actress’ faces, we conducted Bayes factor analyses for repeated-measures ANOVA and paired t-tests. We calculated the Bayes factor in the program R v.3.5.1 (www.r-project.org) using the function anovaBF and ttestBF from the package BayesFactor (Morey and Rouder, 2015). We conducted Bayes factor analyses based on the default priors for ANOVA and paired t-test design (scale r on an effect size of 0.707). A Bayes factor indicates how much more likely each alternative model is supported compared with the null hypothesis.

### Experiment 4: BOP is necessary for modulations of empathic brain activity

#### Stimuli and procedure

These were the same as those in Experiment 3 except the following. Before EEG recording, the participants were informed that all the 16 faces were patients and they were from two baseball teams (half from Tiger team and half from Lion team). After the identity memory task, they performed identity judgments on faces with neutral or pain expressions by pressing one of two buttons while EEG was recorded.

#### EEG data acquisition and analysis

These were the same as those in Experiment 3.

### Experiment 5: Empathic brain activity mediates relationships between BOP and empathy/altruistic behavior

#### Stimuli and procedure

The stimuli and behavioral tests were the same as those in Experiment 1 to assess BOP effects on self-report of perceived pain intensity and altruistic decisions. Thereafter, the participants went through the EEG session that was the same as that in Experiment 3 to examine BOP effects on empathic brain activity. These designs allowed us to test whether BOP induced changes of empathic brain activity plays a mediator role in the pathway from belief changes to altered subjective feelings of others’ pain and altruistic decisions.

#### Behavioral and EEG data recording and analyses

These were the same as those in Experiments 1 and 3.

#### Multiple mediation model analysis

We constructed a serial mediation model to test the hypothesis that BOP (dummy coded as 0 for patients and 1 for actors/actresses) effect on monetary donations was sequentially mediated by two chain mediators, i.e., empathic neural responses and subjective feelings of others’ pain. This model includes three indirect paths: (1) indirect effect of BOP on monetary donation via empathic neural responses (i.e. P2 amplitude); (2) indirect effect of BOP on monetary donation via subjective feelings of others’ pain (pain intensity); (3) indirect effect of BOP on monetary donation via P2 amplitude × pain intensity. To do this, we estimated seven regression models: 1) whether the independent variable (BOP) significantly accounts for the dependent variable (monetary donation) when not considering the mediator (e.g., Path c); 2) whether the independent variable (BOP) significantly accounts for the variance of the presumed mediator (P2 amplitude) (e.g., Path a_1_); 3) whether the independent variable (BOP) significantly accounts for the variance of the presumed mediator (pain intensity) (e.g., Path a_2_); 4) whether the first independent mediator (P2 amplitude) significantly accounts for the variance of the second mediator (pain intensity) (e.g., Path d_21_); 5) whether the presumed mediator (P2 amplitude) significantly accounts for the variance of the dependent variable (monetary donation) when controlling the independent variable (BOP) (e.g., Path b_1_); 6) whether the presumed mediator (pain intensity) significantly accounts for the variance of the dependent variable (monetary donation) when controlling the independent variable (BOP) (e.g., Path b_2_); 7) whether the independent variable (BOP) significantly accounts for the variance of the dependent variable (monetary donation) when controlling the presumed the two mediators (e.g., Path c’). To test the significance of the three paths, we separately conducted to examine the significance of indirect effect (a_1_ × b_1_) of BOP on monetary donation via the P2 amplitude; indirect effect (a_2_ × b_2_) of BOP on monetary donation via pain intensity; indirect effect (a_1_ × d_21_ × b_2_) of BOP on monetary donation via P2 amplitude × pain intensity. Similarly, the bootstrapping method was used to estimate the effect-size and test the hypothesis.

#### Implicit association test

To assure our experimental manipulation of patient and actor/actress identities, after the EEG recording, participants were asked to complete a modified implicit association test (IAT, Greenwald et al., 1998). The participants were asked to respond to faces with patient identifies and patient related words (e.g. ache, weak) with one key and to faces with actor/actress identities and actor/actress related words (e.g. imitation) with another key in two blocks of trials (60 trials in each block). They were then asked to respond to faces with patient identities and actor/actress related words with one key and to faces with actor/actress identities and patient related words with another key in two additional blocks of trials. A D score was then calculated based on response times according to the established algorithm (Greenwald et al., 2003). A positive D score significantly larger than zero would suggest that patient faces were more strongly associated with patient (vs. actor/actress) relevant words whereas actor/actress faces were more strongly associated with actor/actress (vs. patient) relevant words.

### Experiment 6: Neural structures underlying BOP effects on empathy

#### Stimuli and procedure

We adopted 24 video clips from 6 models from our previous work (Luo et al., 2014) and recorded 56 video clips from 14 Chinese models (half males) in Experiment 6. Each model contributed four video clips, in which a face with pain expressions receiving painful stimulation (needle penetration) or with neutral expressions receiving non-painful stimulation (cotton swab touch) applied to the left or right cheeks. Each video subtended a visual angle of 21° × 17° (width × height) at a viewing distance of 80 cm during fMRI scanning.

A photo of each model with a neutral expression was obtained from each video clip. These photos were then used in the identity memory task, which was the same as that in Experiment 3. After the identity memory task the participants underwent fMRI scanning. An event-related design was employed in 6 functional scans. Each scan consisted of 20 video clips (half patients (5 pain and 5 neutral expressions) and half actors/actresses (5 pain and 5 neutral expressions)) that were presented in a random order. Each video clip lasted for 3 s. There was a 9-s interstimulus interval between two successive video clips when the participants fixated at a central cross and had to judge the identity (patient or actor/actress) of each model in the video clip by pressing one of two buttons using the right index or middle finger. The relation between responding finger and face identity was counterbalanced across participants.

After fMRI scanning, the participants were presented with each video clip again outside the scanner. They were asked to rate pain intensity of each model (1 = not painful at all; 7 = extremely painful) and own unpleasantness (1 = not unpleasant at all, 7 = extremely unpleasant). Finally, we assessed the participants’ beliefs of models’ identities by asking them to answer the question of “To what extent do you believe the identity of this model (either patient or actor/actress)?” on a 15-point Likert-type scale (−7 = extremely believed to be an actor/actress, 0 = not sure, 7 = extremely believed to be a patient).

#### fMRI data acquisition and analysis

Imaging data were acquired using a 3.0 T Siemens scanner with a standard head coil. Head motion was controlled to the maximum extent by using foam padding. Functional images were acquired by using T2-weighted, gradient-echo, echo-planar imaging (EPI) sequences sensitive to Siemens scanner contrast (64×64×32 matrix with 3.75×3.75×5 mm^3^ spatial resolution, repetition time = 2000 ms, echo time = 30 ms, flip angle = 90°, field of view = 24×24 cm). Anatomical images were subsequently obtained using a standard 3D T1-weighted sequence (256×256×144 matrix with a spatial resolution of 1×1×1.33 mm3, TR = 2530 ms, TE = 3.37 ms, inversion time (TI) = 1100 ms, FA = 7°).

Functional images were preprocessed using SPM12 software (the Wellcome Trust Centre for Neuroimaging, London, UK, http://www.fil.ion.ucl.ac.uk/spm). Functional scans were first corrected for within-scan acquisition time differences between slices and then realigned to the first volume to correct for inter-scan head motions. This realigning step provided a record of head motions within each fMRI run. Head movements were corrected within each run and six movement parameters (translation; x, y, z and rotation; pitch, roll, yaw) were extracted for further analysis in the statistical model. The functional images were resampled to 3 × 3 × 3 mm^3^ voxels, normalized to the MNI space using the parameters of anatomical normalization and then spatially smoothed using an isotropic of 8 mm full-width half-maximum (FWHM) Gaussian kernel.

Whole-brain analyses was conducted to examine brain regions in which activities increased in response to pain vs. neutral stimuli regardless of patient or actor/actress identities. This contrast pooled video clips of patient and actor/actress models together to focus on BOLD responses to painful vs. neutral stimuli. The design matrix also included the realignment parameters to account for any residual movement-related effect. A box-car function was used to convolve with the canonical hemodynamic response in each condition. Random-effect analyses were conducted based on statistical parameter maps from each participant to allow population inference. The contrast values were compared using whole-brain paired t-tests to identify activations, which were defined using a threshold of voxel-level p < 0.001, uncorrected, cluster-level p < 0.05, FWE corrected. We also conducted a whole-brain analysis to calculate the contrast of patient vs. actor/actress non-painful stimuli to test whether BOP may motivate inference of patients’ mental states independently of any perceived painful cues.

#### Representational similarity analysis

We conducted a representational similarity analysis (RSA) of brain activity (Nili et al., 2014) to examine neural correlates to BOP effects on subjective feeling of others’ pain. We constructed a 4 × 4 dissimilarity matrix (DM) for each participant with each cell in the DM represents the mean difference in rating scores of pain intensity between each pair of conditions. The DM was then used for a whole-brain searchlight RSA to identify brain regions in which the pairwise similarity of neural responses in the 4 conditions (2 Expressions × 2 Identities) corresponded to the behavioral DM of condition dissimilarity in each participant. To do this, functional images were similarly preprocessed using a GLM but were not smoothed and normalized. We then estimated a GLM for each participant with Identity (Patient vs. Actor/Actress) and Expression (Pain vs. Neutral) as experimental regressors. The estimated beta images corresponding to each condition were then averaged across runs at each voxel and were used as activity patterns in the RSA toolbox (Nili et al., 2014). We compared the neural-pattern similarity (i.e., the neural DM) with the behavioral DM in each voxel of the brain using the searchlight procedure (Kriegeskorte et al., 2006). The neural DM was constructed by 1 minus the correlation coefficient between the pattern vectors of each condition pair. The Spearman rank correlations between the neural DM and behavioral DMs were computed and assigned to the central voxel of the sphere. As such, the searchlight procedure produced Spearman p values on each voxel for each participant, which were then subject to Fisher’s z transformation for statistical tests. The resulting z maps were then normalized to standard space (resampled to 3 x 3 x 3 mm^3^ voxels), smoothed (FWHM= 8mm), and entered into a random effect analysis using one-sample t tests against zero. Significant results were reported using a threshold of P (FWE) < 0.05 at the voxel level.

## Data availability

The data that support the findings of this study are available from the corresponding author upon reasonable request.

## Code availability

The code used to analyse the data that support the findings of this study are available from the corresponding author upon reasonable request.

## Acknowledgements

This work was supported by the Ministry of Science and Technology of China (2019YFA0707103) and the National Natural Science Foundation of China (projects 31871134, 31421003, and 31661143039). The authors thank T. Gao, X. Han, T. Huo, S. Mei, C. Pang, Y. Pu, X. Wang, G. Zheng, N. Zhou, Y. Zhou for proofreading the manuscript. The funder had no role in the conceptualization, design, data collection, analysis, decision to publish or preparation of the manuscript.

## Author contributions

S.H. and T.W. conceived the research programme and designed the experiments. T.W. and S.H. carried out the experiments and analysed the data. S.H. and T. W. wrote the paper. S.H. supervised the entire work.

